# The addition of antibiotics to embryo culture media caused altered expression of genes in pathways governing DNA integrity in mouse blastocysts

**DOI:** 10.1101/2022.03.29.486218

**Authors:** Qianqian Han, Yan Li, Xu Ji, Lu Chang, Wenjuan Li, Jianfeng Shi, Jing Liu, Wuhua Ni, Xuefeng Huang, Chris O’Neill, Xingliang Jin

**Affiliations:** Reproductive Medicine Center, The First Affiliated Hospital of Wenzhou Medical University, Wenzhou, Zhejiang Province, China, 325000; National Institutes for Food and Drug Control of China, No.31 Huatuo road, Daxing district, Beijing, China 102629; School of Pharmaceutical Sciences, Nanjing Tech University, Nanjing, Jiangsu Province, China, 211816; Sydney Center for Regenerative and Developmental Medicine, Kolling Institute for Medical Research, Sydney Medical School, University of Sydney, St. Leonards, New South Wales, Australia, 2065

**Keywords:** Antibiotic, differential gene expression, DNA recombination, embryo development, gene otology, in vitro fertilization, signal pathway, tumorigenic

## Abstract

Antibiotics are common components of embryo culture media and minimize the risk of microbial contamination and infection during assisted reproductive technology procedures (ART). The effects of two aminoglycoside antibiotics (gentamicin, streptomycin) and penicillin on the global profiles of gene expression (DE) were assessed by RNA-seq of individual mouse blastocysts. Zygotes were cultured in an optimized defined medium formulation (KSOM) to which a dose range of each antibiotic was added. A dose-dependent retardation of the rate of zygote development to morphologically normal blastocyst was observed and this was accompanied by a reduction in the number of cells present within the resulting blastocysts. The lowest dose of each antibiotic tested (similar to the concentrations used in clinical grade media) caused significant differential expression of approximately 1800 genes. In most cases antibiotic treatment caused a reduction in gene expression and gene ontology analysis showed that down regulated genes were enriched for several biological processes related to the maintenance of genomic integrity. All three antibiotics caused the downregulation of *Brca2, Blm, Rad51c* and *Rad54l*, genes involved DNA homologous recombination pathways and also several p53-dependent genes. Immunolocalization studies showed that each antibiotic also reduced level of BRCA2 and RAD51C detected within blastocysts. The present study shows that the supplementing embryo culture media with antibiotics is associated with wide ranging alterations in gene expression in a manner that could potentially compromise the genomic integrity of the resulting embryos.

## Introduction

The creation of embryos by in vitro fertilization and embryonic culture procedures is an important treatment of infertility in humans. Is generally recognized that the resulting embryos have a lower viability and developmental potential than embryos conceived naturally. There have been many developments in embryo culture techniques, including improved media formulations(1, 2), yet the addition of antibiotics remains a common component of these procedures and serves as a quality assurance against microbial infection of cultures in vitro. The types of antibiotics used and their concentration varies between media formulations. The most commonly used antibiotics are penicillin, streptomycin or gentamicin(3-5).

The aminoglycoside antibiotics (streptomycin or gentamicin) display concentration-dependent bactericidal activity against “most gram-negative aerobic and facultative anaerobic bacilli” but not against gram-negative anaerobes and most gram-positive bacteria(6). They act to cause codon misreading by binding to the 30S ribosomal subunit and blocking the translocation of peptidyl-tRNA from the acceptor site to the donor site(7, 8). Penicillin antibiotics may be effective against many bacterial infections caused by staphylococci and streptococci. Several studies have reported adverse effects of the addition of antibiotics to media upon the rates of embryonic development in vitro in human(9, 10), hamster(11) and mouse(4, 12), but the underlying causes of this effect have yet to be studied in detail.

This study analyzed the influence of two aminoglycosides and penicillin in the mouse zygote development and then subjected the resulting blastocysts to global gene expression analysis by single embryo RNA-seq. The results show that at the dose commonly used in clinical grade media each class of antibiotic caused pervasive changes in the pattern of gene expression. The predominant effect was for a downregulation of gene expression, and this occurred across a range of regulatory pathways, including those governing the integrity of the genome.

## Materials and Methods

### Animal experiments

Animal experiments were conducted with the approval by the ethics committee of Wenzhou medical university. Hybrid (C57BL/6 X CBA/He) mice in were housed and bred in Wenzhou Medical University. Ovulation was induced in six-week-old female mice by intraperitoneal injection of 5 IU pregnant mare serum gonadotropin (PMSG) (Ningbo Second Hormone Factory, China). After 48 h, mice were injected with 5 IU human chorionic gonadotrophin (hCG) (Livzon, Zhuhai, China). Pregnancy was confirmed by the presence of a copulation plug the following morning. Zygotes were recovered 18 h post-hCG from mated females in HEPES-buffered human tubal fluid medium (HEPES-HTF) (13). Cumulus cells were removed by brief exposure to 300 IU hyaluronidase (Sigma Chemical Company, St Louis, MO, USA). Embryos were cultured at density of 10 embryos in 10 µl KSOM medium supplemented with 3 mg bovine serum albumin/mL (14) in 60-well plates (LUX 5260, Nunc, Naperville, IL, USA) overlaid with 2 mm of heavy paraffin oil (Sigma) at 37°C in 5% CO_2_ in air. All components of KSOM medium were embryo culture grade (Sigma). The working concentrations of three antibiotics (gentamicin sulfate salt (Sigma G-1264), penicillin-G sodium salt (Sigma-P3032, ≥1477U/mg) and streptomycin sulfate (Sigma S-9137, ≥ 720 IU/mg)) were prepared by dissolving directly in KSOM medium on the day of use. The treatments were then prepared by a serial dilution in KSOM to produce: (i) control medium without antibiotics; (ii) 0.01, 0.1, 1 mg Gentamicin/mL; (iii) 72, 720, 7200 IU Penicillin/mL; (iv) 0.05, 0.5, 5 mg Streptomycin/mL. Zygotes were cultured for 96 h to assess their effects on developmental outcomes and gene expression.

### Single blastocyst RNA-seq

Six of individual morphologically normal blastocysts from 3 replicates of each treatment with the lowest dose of antibiotics was selected and subjected to RNA-seq. RNA Seq cDNA preparation was as described, with modification (15). Briefly, each blastocyst was washed in cold PBS and then lysed in the buffer containing 10 X PCR buffer II, 25mM MgCl2, 10% NP40, 0.1M DTT, 0.5 µM UP1, 2.5mM dNTPs, SUPERase• In™ RNase Inhibitor and Ambion™ RNase Inhibitor at 70°C for 90s to obtain whole RNA. RNA was reverse transcribed with SuperScript III RT and T4 gene 32 protein to synthesize the first strand cDNAs. After free primers were removal by Exonuclease I, the 3’ end of the first stranded cDNA was added with poly(dA) tail by the reaction with RNase H, 100 mM dAMP, terminal deoxynucleotidyl transferase. Second strand cDNAs were synthesized by dNTPs, AUP2 primer, TakaRa EX Taq HS and they were 5’-UP2-(T)n-cDNA-(A)n-UP1-3’. The first amplification of cDNAs was performed with TakaRa EX Taq HS and UP1 primer by 20 cycles. The cDNAs were purified with QIAQuick PCR purification kit and then amplified by second round of 9 cycles PCR with Amine-blocked UP1, amine-blocked UP2 and TakaRa EX Taq HS. After purification of the cDNAs with QIAQuick PCR purification kit, the concentration of cDNAs was measured with Qubit. Approximately 1 ng of this cDNA was processed with TruePrep DNA library Prep Kit V2 for Illumina (Vazyme, Nanjing, China), according to the manufacturer’s instructions. After the fragmentation step of the full-length transcripts, a 10-cycle PCR amplification was performed using a unique combination of index 1 (i7) and index 2 (i5) adapters per library. The average length distribution of the fragmented cDNA libraries was assessed by the high sensitivity DNA kit (Agilent, USA) and the libraries quantified by Qubit (Thermo Scientific, USA), prior to equimolar pooling. Libraries were sequenced using Illumina HiSeq X Ten (Illumina, San Diego, CA, USA). All reagents were bought from Thermo Fisher Scientific unless mentioned specifically. The cDNA library reads generated are deposited at https://dataview.ncbi.nlm.nih.gov with the BioProject accession number PRJNA820513.

The raw sequencing reads (FASTQ files) were trimmed with Trimmomatic (V0.38) to remove the adapters, discard low-quality reads and obtain the clean reads (16). The reads were aligned to the reference genome (Mus_musculus. GRCm38) to obtain bam files using HISAT2 (V2.1.0) (17). FPKM (fragments per kilobase of exon model per million mapped reads) and TPM (transcripts per million) were used for With-sample Normalization. Using StringTie (V1.3.4d) the alignment files were reconstructed and the transcripts were assembled. After initial assembly, the assembled transcripts are merged together, which creates a uniform set of transcripts for all samples (18). The expression levels of each gene and transcript were quantitated and the differences in expression for all genes among the different experimental conditions was calculated (18, 19). Transcript_ratio and Gene_ratio was calculated. QualiMap (V2.2.1) was used for the statistical analysis of 5’bias, 3’bias and reads_exonic_ratio, and RSeQC (V3.0.0) for rRNA_ratio. Differential gene expression between no antibiotic control and each antibiotic treatment was analyzed using the R statistical package DESeq2 (V1.22.2) (20). Bioinformatic analysis of the gene ontology (GO) analysis of differentially expressed genes was performed using the Enrichr tool (http://amp.pharm.mssm.edu/Enrichr) (21). On ontologies catalog, the analyzed GO terms included Biological Process 2018 and KEGG 2019 Mouse Pathways (Kyoto Encyclopedia of Genes and Genomes).

### Quantitative Reverse transcription – PCR (qRT-PCR)

qRT-PCR was performed as described previously(22). Individual blastocysts derived from embryo culture was washed in cold PBS 3 times to remove the media and transferred in minimal volume to 2 µL of RNA buffer containing 0.1 IU RNase inhibitor (Thermo Fisher Scientific). RNA was extracted by three repeats of freezing in liquid nitrogen and thawing with vortex. The RNA was purified with RQ1 RNase-Free DNase Kit (Thermo Fisher). cDNA synthesized was performed with 1^st^ strand cDNA kit (Thermo Fisher) according to manufactory instruction. Negative controls were either reactions without reverse transcriptase or without the RNA sample replaced by DEPC treated MilliQ water (to test for any RNA or DNA contamination). An internal positive control was to test for expression of *Actb*. Amplification of cDNA was performed with sequence specific primers as follows (Table 1) and SYBR Green PCR master mix (Thermo Fisher) on Step One Plus Real-Time PCR System (Thermo Fisher). *Ct* values were calculated by the system software. The Delta *Ct* was a measure of relative changes in the transcripts of the tested gene content of the embryo to housekeeper gene *Actb*. A plot of 2^*-(normalized DeltaCt)*^ was shown.

**Table 1.**
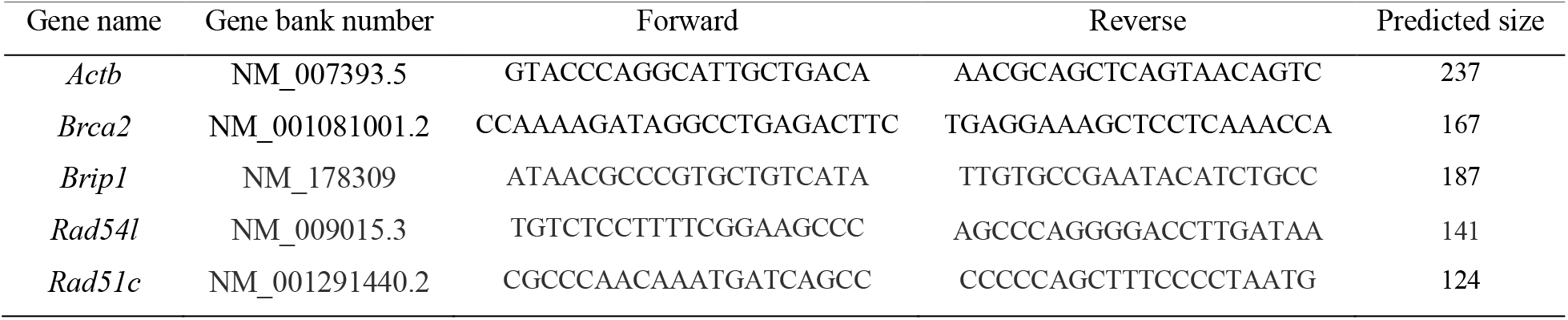
The primer sequences for RT-PCR analysis of targeted genes and reference gene.

The PCR products were analysed on 3% agarose gel electrophoresis. Samples of each transcript were sequenced to confirm identity.

### Immunofluorescence

Immunofluorescence was performed as previously described (23, 24). After fixation, permeabilization and blocking, embryos were incubated overnight at 4°C with primary antibodies: 2 μg/mL rabbit anti-NANOG polyclonal IgG, 2 μg/mL rabbit anti-BRCA2 polyclonal IgG, 2 μg/mL mouse anti-RAD51C polyclonal IgG and 2 μg/mL isotype negative control immunoglobulin. All primary antibodies were purchased from Abcam (Cambridge, MA, USA). Primary antibodies were detected by Texas Red-conjugated goat-anti mouse (Sigma) or FITC-conjugated goat anti-rabbit (Sigma) secondary antibodies for 1 h at room temperature. Nuclear localization was determined by encountered staining with 10 μg/mL Hoechst33342 (Sigma) or 0.1µg/mL propidium iodide (PI). Whole section imaging was captured by Nicon microscope with x 20 len. All quantitative analysis of immunofluorescence experiments was performed with Image-Pro Plus (version 6.3, Media Cybernetics USA). The number of the inner cell mass (ICM) was determined by counting the cells that expressed NANOG, and total cell number by counting of Hoechst33342 stained nuclei in the blastocyst.

### Statistical Analysis

Statistical analyses of differences in development outcomes, RT-PCR results or staining levels were performed with SPSS for Windows (Version 22.0, SPSS Inc., Chicago, IL, USA). Normalized *Ct* value and cell number were quantitatively analyzed by univariate analysis of variance. Those parameters were set as the dependent variable, while the antibiotic treatments were the independent variables. Experimental replicates were incorporated into the model as covariates. Differences between individual treatments were analyzed by the least significance difference test. Blastocyst development rate was assessed by binary logistic regression analysis.

## Results

### Antibiotics caused the adverse developmental viability in vitro

The zygotes were cultured in the KSOM medium supplemented with a range of 3 doses of each antibiotic. Each treatment caused a dose-dependent reduction in the proportion of zygotes developing to morphological blastocysts (Figure 1 A, B, C), and also the total number of cells and the cells within the inner cell mass in the resulting embryos (Figure 1 D, E, F).

**Figure 1.**
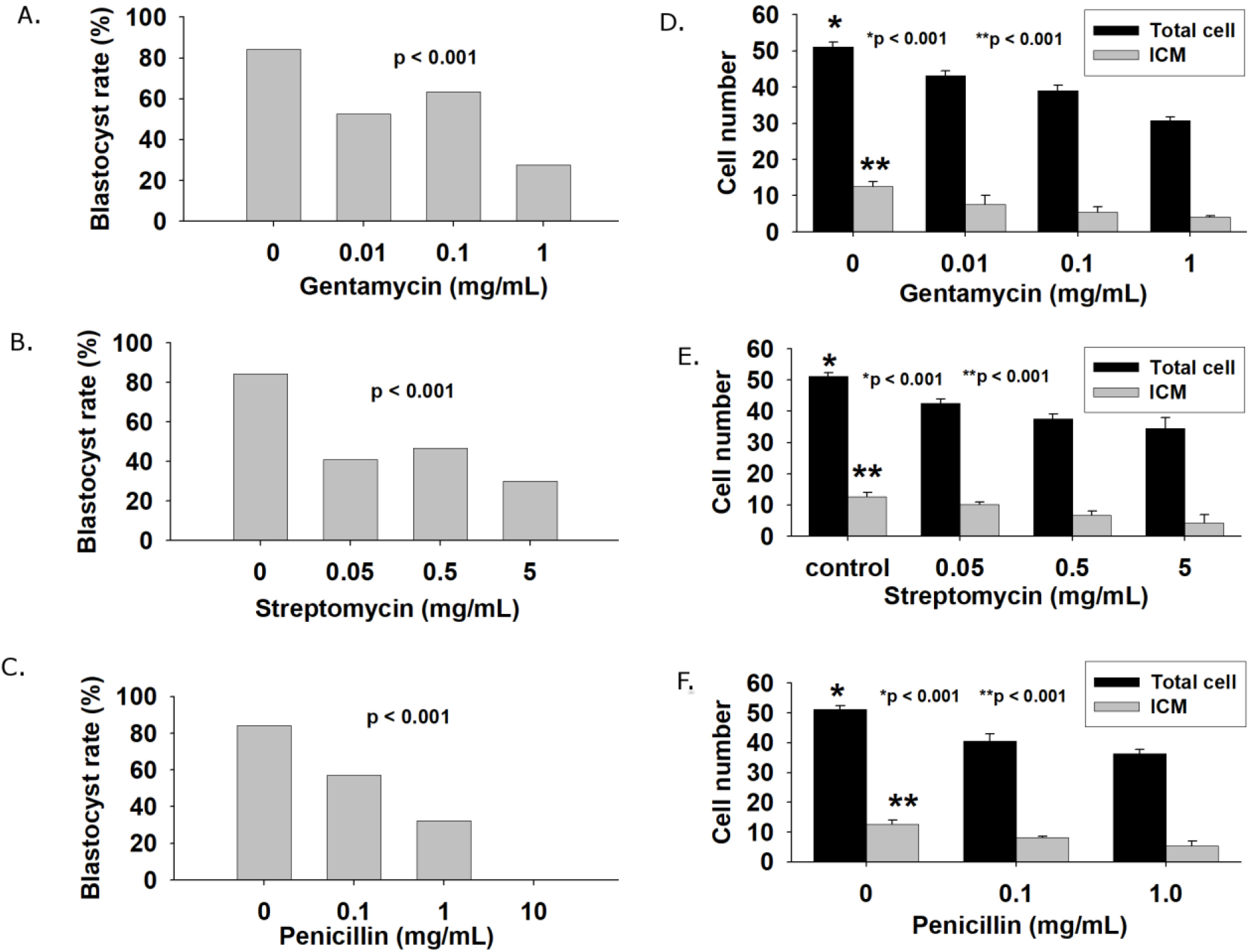
the effect of antibiotics on mouse zygote development in vitro The blastocyst formation rate and the number of total cell and ICM in the blastocysts derived the KSOM medium supplemented with addition of gentamycin (A, D), streptomycin (B, E) and penicillin (C, F). The results were representative of three independent replicates, and each replicate had at least 20 embryos for each dose of antibiotic. p < 0.001, dose-dependent effect on blastocyst formation rates between the control (without antibiotics) and a range of antibiotic doses. * p < 0.001, total number of cells in the blastocyst compared to each dose of the antibiotic. **p < 0.001, the number of ICM cells compared to each dose of the antibiotic.

### RNA-seq analysis of antibiotics-related differential expression of transcriptome in the blastocysts

Representative morphologically normal blastocysts from the control and the lowest dose of each antibiotic treatment were selected and subjected to individual embryo RNA-Seq analysis. The cDNA produced from each replicate sample was not significantly different between four treatment groups (Suppl 1). RNA-Seq produced more than 23 million reads the library and the counts were not different between each replicate sample (Suppl Table 1, Figure 2-A, B), of which more than 85% could be successfully aligned to the reference mouse genome (Suppl Table 1). The commonly expected duplicates rates were shown(25). A high ratio of exonic_reads was obtained and almost no rRNA remained in treatment samples (Suppl Table 1), indicating a satisfactory quality of the poly(A) RNA-seq libraries(26, 27). By comparison, the data for negative control (No-RT) produced a low output of cDNA and reads. The data for a range of -5 <log2-fold-change < 5 that differed between negative control and no-antibiotic control were considered as noise distribution and ignored for further analysis (Suppl 1), and thus, the data for 5 ≤log2-fold-change≤ -5 with adjusted p-value <0.05 were used for the analysis of differential expression between the control and antibiotics treatments.

**Figure 2.**
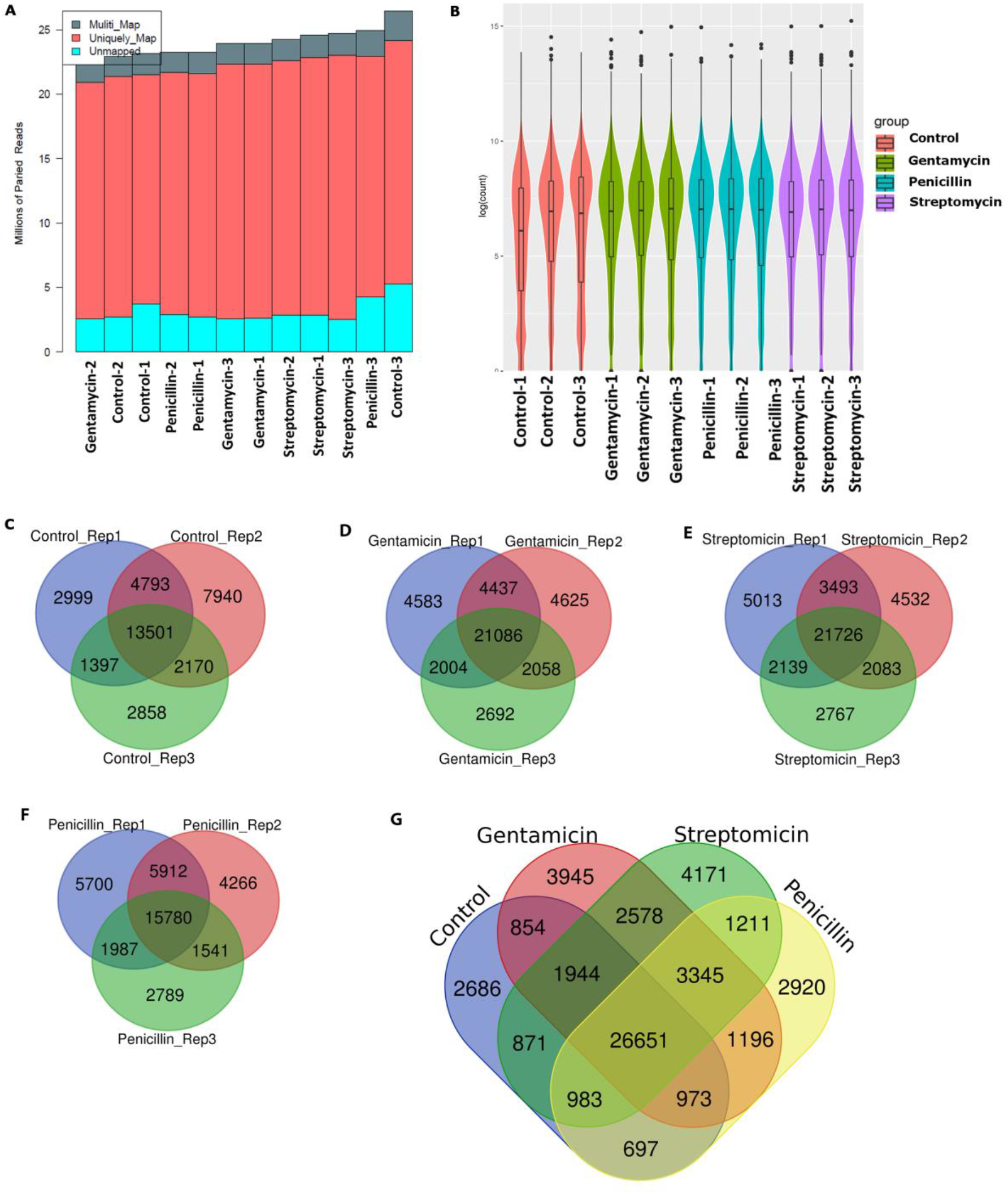
RNA-seq analysis of antibiotic-caused DET in single blastocyst. (A) A bar plot showing the number of paired reads to reference transcriptome for each replicate sample; (B) Violin plot showing the Log counts measured to each replicate sample (y-axis: Log10-scaled) across replicates from four treatment groups (x-axis). Violins were grouped and colored by distinct treatment groups of library preparation. Venn diagram illustrating the relationship and overlap of transcripts identified by RNA-seq in the blastocysts derived from the replicates of same treatment (C); and the antibiotics and control groups (D). Venn diagram was colored by distinct antibiotics and control treatments.

Venn diagram analysis demonstrated the more than 47% similarity of transcripts between the replicates of same treatment (Fig 2-C, D, E, F), and the 26651 transcripts were commonly detected between 4 treatment groups (Figure 2-F) while markedly few unique transcripts were found in each treatment group. Comparison of global gene expression in the blastocysts between control and each antibiotic treatment showed the differential expression of ∼1800 genes accounted for transcriptome change, of which the greater part of genes was down regulated (Figure 3 A-C, Suppl 2, 3, 4). Gentamicin and streptomycin gave more downregulated genes than penicillin.

**Figure 3.**
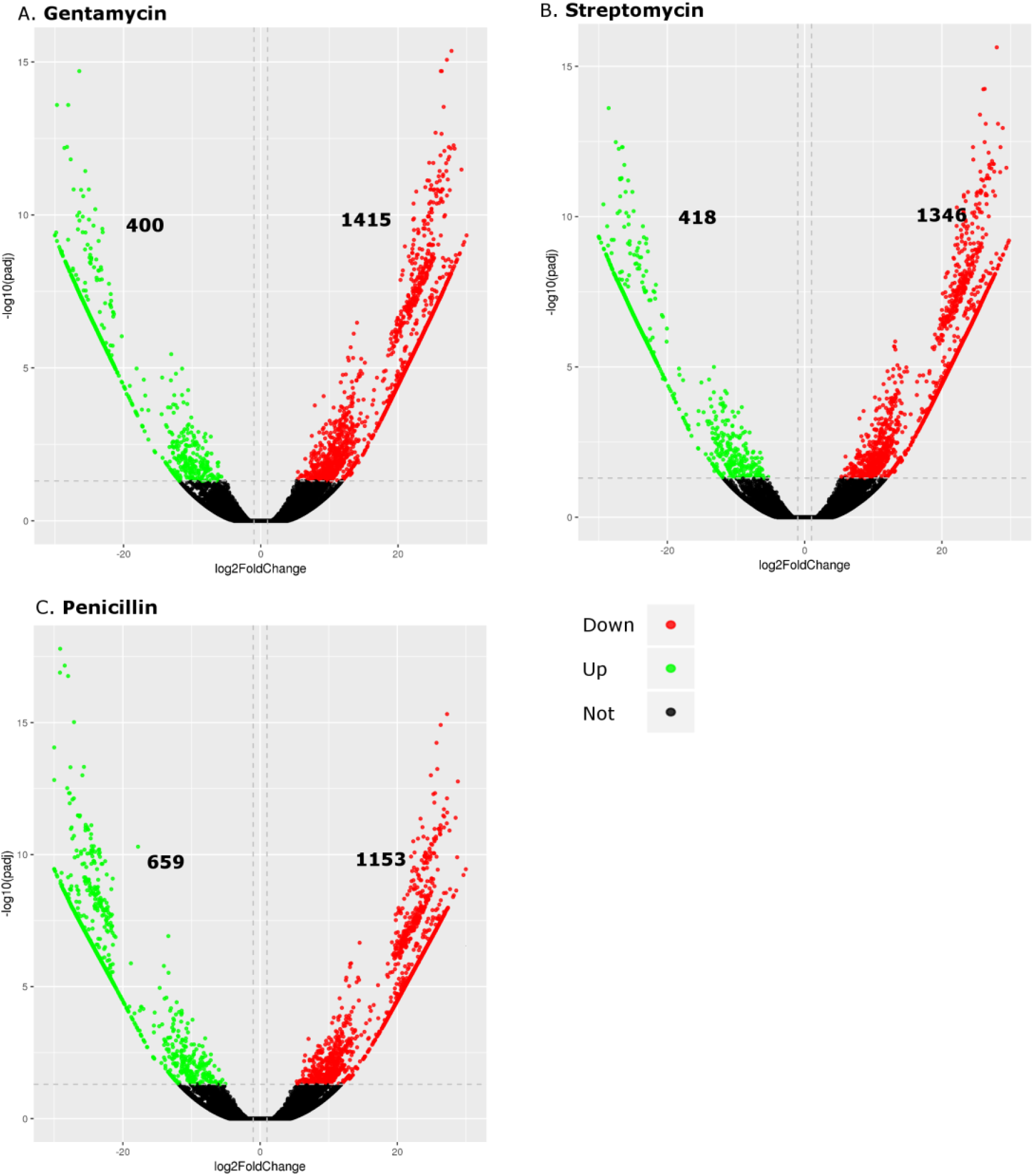
The effect of three antibiotics on transcription compared to no treatment control in the blastocyst. The Volcano plot showed the differentially expressed genes according to the p-adj and log2Fold change values. Distinct colors presented the down- and up-regulated, and unregulated transcripts, and the number of transcripts was shown.

### Antibiotics modulated a range biological processes and KEGG pathways associated with maintenance of genomic integrity

Genes downregulated as a result of gentamicin treatment were associated with the enrichment of 72 GO biological process (BP) and the top 10 included several DNA biological processes, amyloid precursor protein metabolism, protein modification and cell cycle (Figure 4 A-a, Suppl Table 2, Suppl 5). Top three BP terms were involved in DNA biological roles, including: DNA metabolic process, DNA recombination and cellular response to DNA damage stimulus (Suppl 5). Gentamicin treatment impacted P-value ranked 21 KEGG pathways with the top 10 listed including the cell cycle, mannose type O-glycan biosynthesis, RNA transport, homologous recombination pathways (Figure 4 A-b, Suppl Table 3, Suppl 5).

**Figure 4.**
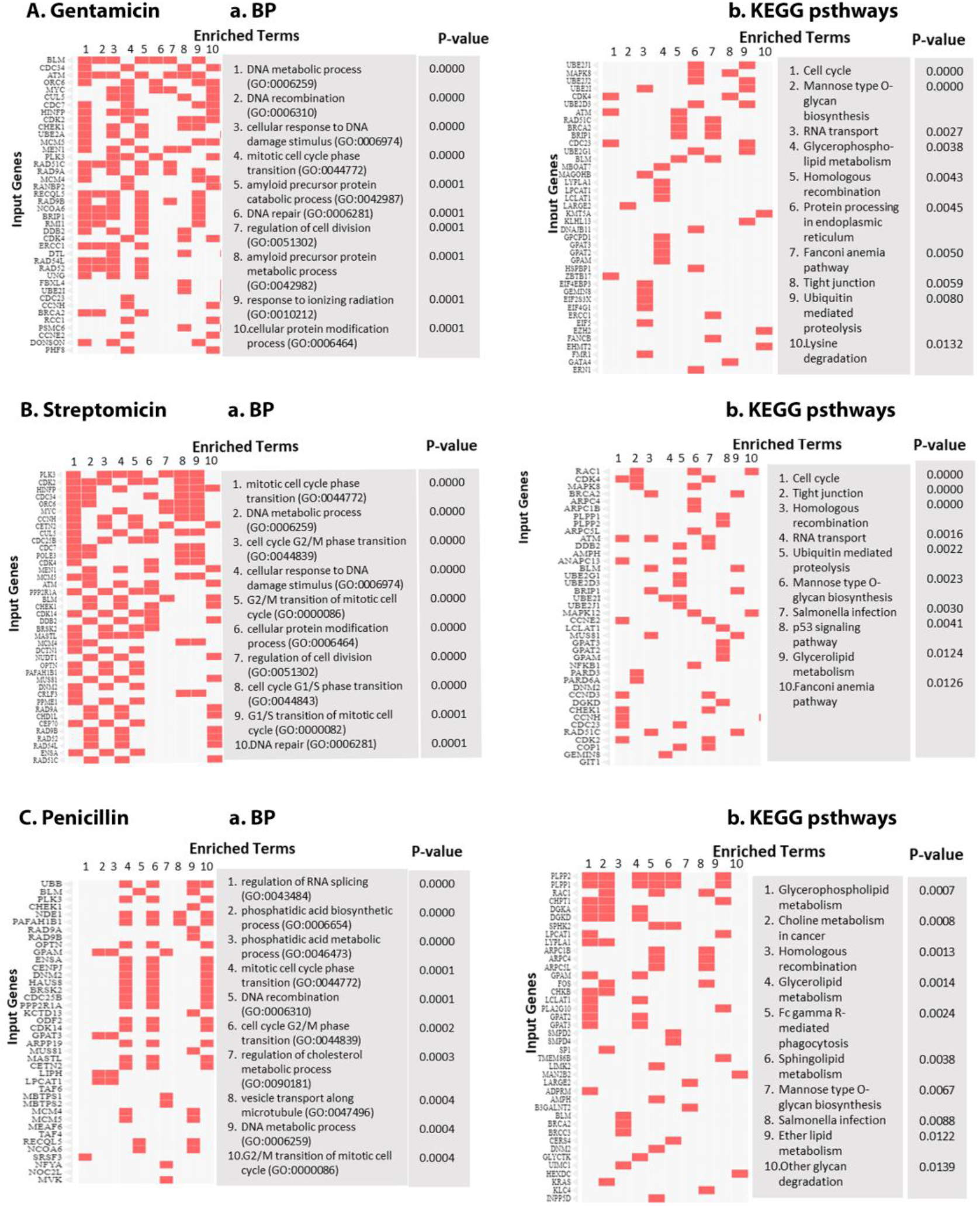
The effects of antibiotics-downregulated genes on the biological function, relevant signal pathways and prospective health conditions. P-Value ranked clustergrams showed that gentamicin (A), Streptomicin (B) and penicillin(C) on the association of the genes with top 10 of the BP and KEGG signal pathway.

Genes down regulated in response to streptomycin involved 66 BP (Suppl 6) and top 10 mainly involved cell cycle and DNA biological processes, including mitotic cell cycle phase transition, DNA metabolic process and cell cycle G2/M phase transition (Figure 4 B-a, Suppl Table 2, Suppl 6). There were 27 KEGG signal pathways were affected, including cell cycle, Tight junction, Homologous recombination, RNA transport, Ubiquitin mediated proteolysis, Mannose type O-glycan biosynthesis, Glycerolipid metabolism and Fanconi anemia pathway (Figure 4 B, Suppl Table 3, Suppl 6).

Penicillin downregulated genes affected 51 enriched terms of BP (Suppl 7). The top 10 of these included RNA processing, phosphatidic acid biological processing, DNA recombination, cholesterol metabolic process, cell cycle and vesicle transport along microtubule (Figure 4 C-a, Suppl Table 2, Suppl 7). 15 KEGG signal pathways were affected, and three of the top 10 KEGG pathways included Glycerophospholipid metabolism, Choline metabolism in cancer, Homologous recombination, glycerolipid metabolism and Mannose type O-glycan biosynthesis (Figure 4 C-b, Suppl Table 3, Suppl 7).

It was noted that there were some similarities on the impacts of all three antibiotic treatments, included two BP enriched terms: DNA metabolic process (GO:0006259) and mitotic cell cycle phase transition (GO:0044772) (Suppl Table 2) and two KEGG pathways: “Homologous recombination and, Mannose type O-glycan biosynthesis (Suppl Table 3). These two KEGG pathways involved two groups of genes: *Rad52, Blm, Rad51c, Rad54l, Brcc3, Brca2; B3galnt2, Pomt1, Pomgnt2, Pomgnt1, Large2*, respectively. The key differences between the three antibiotics were that *Atm* was also involved in gentamicin and streptomycin-down-regulated “Homologous recombination”, and *Uimc1* in penicillin (Suppl Table 3).

It is noted that antibiotics caused down-regulation of transcription of a range *Trp53*-dependent BP enriched terms. All three affected “regulation of signal transduction by p53 class mediator (GO:1901796)”, which involved in a range of genes including *Plk3, Blm, Meaf6, Taf9, Noc2L, Hipk1, Aurkb, Tpx2, Ubb, Chek1, Taf6, Kmt5a, Rbbp7, Rad9b, Taf4, Rad9a*. (Suppl Table 4). The key differences were that gentamycin and streptomycin also downregulated *Ehmt2, Brip1, Cdk2* and *Atm*, while Penicillin *Prkab2*.

There were fewer genes up-regulated in response to each of the antibiotics. GO term enrichment analysis of these showed 192 BP terms were affected by gentamicin and top 10 was noted to regulate the roles of protein and ion on the membrane and phosphorylation (Suppl 8, Figure 5 A-a). 33 KEGG signal pathways were found and the most striking was insulin signal pathway that associated with *Raf1, Hras, Phka1* and *Phka2* (Suppl 8 Figure 5 A-b). Streptomycin upregulated genes modulated 129 enriched terms of BP including regulation of defense to virus associated with *Ppm1b, Pcbp2* and *Itch* genes (Suppl 9, Figure 5 B-a), while the genes related to their top 10 BP terms were neither clearly associated with streptomycin-resulted 4 KEGG pathways (Supplemental 9, Figure 5 B-b). 134 BP terms were upregulated by penicillin and 8 of top 10 were involved in the several RNA processing terms (Suppl 10, Figure 5 C-a). They markedly modulated the splicesome KEGG pathway related to upregulated 11 genes (e.g *Ddx5*) (Suppl 10, Figure 5 C-b).

**Figure 5.**
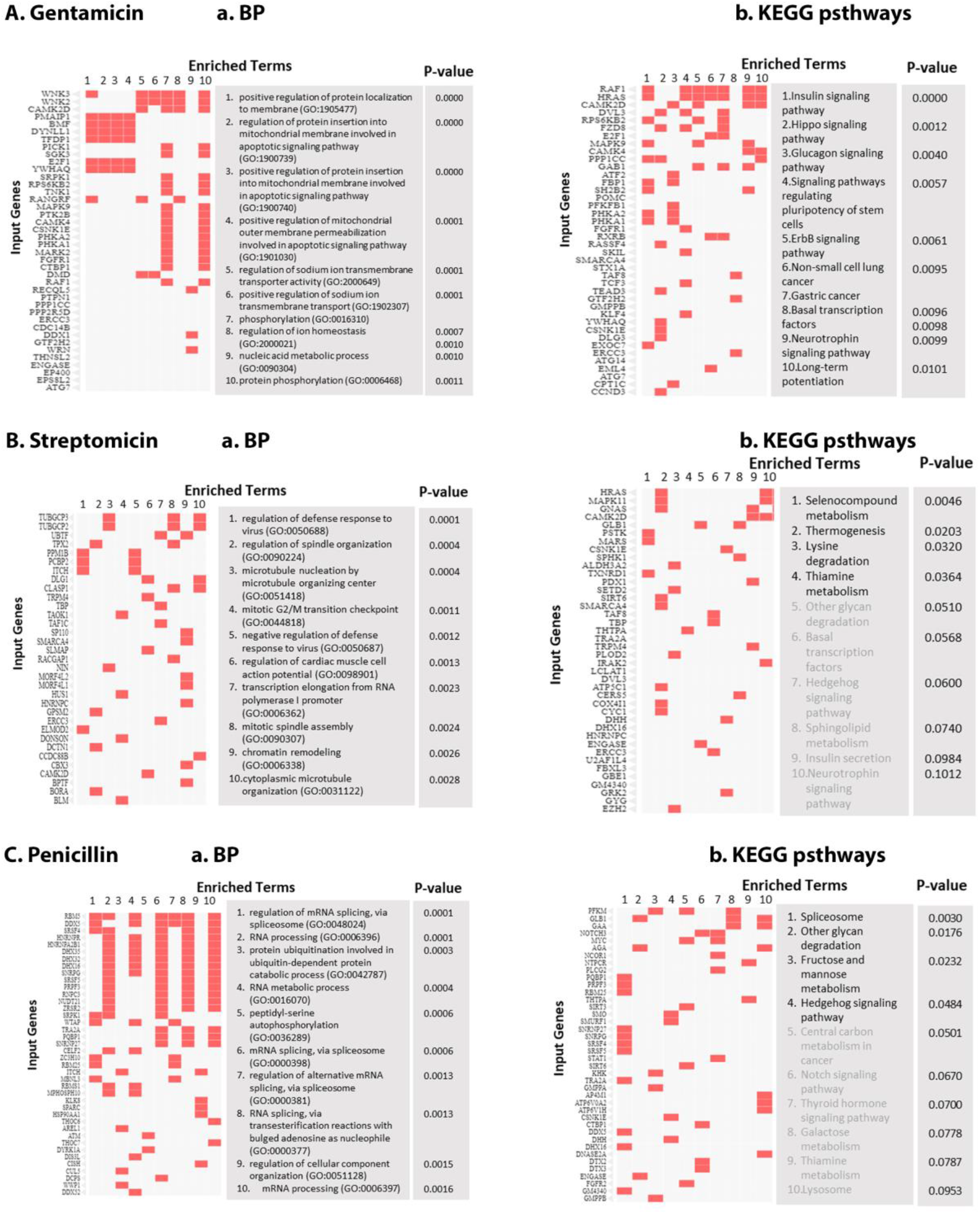
The effects of antibiotics-upregulated genes on the biological function, relevant signal pathways and prospective health conditions. P-Value ranked clustergrams showed that gentamicin (A), Streptomicin (B) and penicillin(C) on the association of the genes with top 10 of the BP and KEGG signal enriched terms, and these were paralleled with the relevant p-values.

### qRT-PCR and immunostaining down-regulated genes in the blastocyst

Quantitative real-time RT-PCR was used to confirm and validate the differential expression of a range of genes detected in the RNA Seq analysis. The reduced expression of *Brca2, Rad51c, Rad54l* and *Brip1* transcripts in the blastocyst that was produced from culture in antibiotics with the lowest dose tested, compared to the control (p < 0.01) (Figure 6 A), excepts *Brip1* was not affected by penicillin. The results were consistent with the finding from RNA-seq analysis. Immunostaining showed that BRCA2 protein was detected and more accumulated in the cytoplasm of the blastocyst derived in no-antibiotic medium (Figure 6 B). Overall, lower level of BRCA2 staining was visualized in the blastocysts from the treatments of three antibiotic and its localization was not characteristic of cytoplasmic accumulation but some in the nuclei as well (Figure 6 B). RAD51C protein was present in the blastocyst with strong dotted staining in the cytoplasm of the trophectoderm cells and relatively even distribution in the inner cell mass cells (Figure 6 C). This cytoplasmic staining was less present in the blastocysts from the treatments of three antibiotics (Figure 6 C), and overall level of the protein was also reduced.

**Figure 6.**
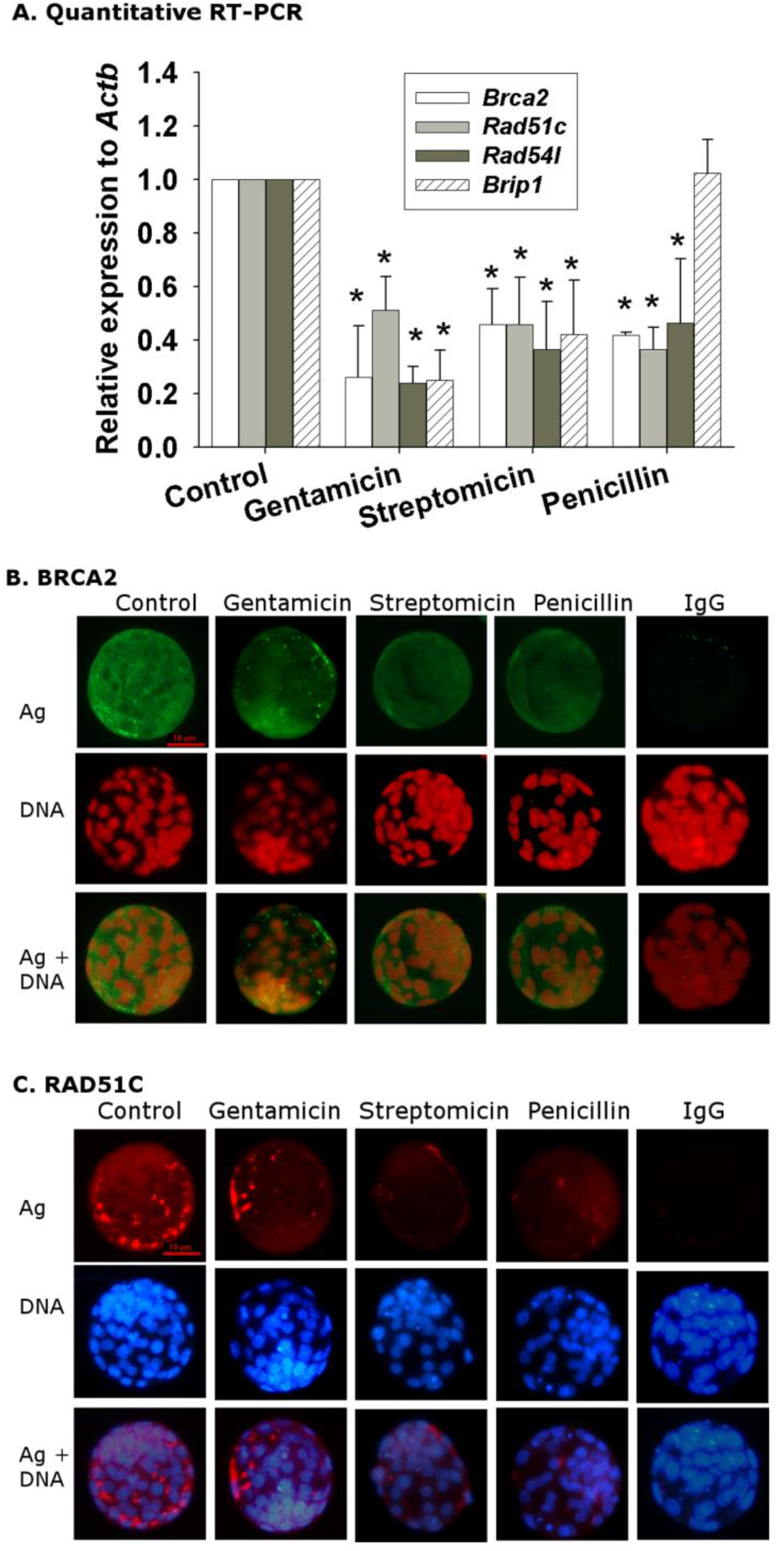
Transcripts and proteins of antibiotics-mediated downregulated genes in the blastocyst (A) qRT-PCR detected the antibiotics-caused reduction in transcription of *Brca2, Rad51c, Rad54l* and *Brip1* in the mouse in vitro-developed blastocyst. The data were representative of the Mean ± STD of normalized ΔΔCt to internal housekeep gene and non-antibiotics treatment (control) for three independent experiments. *P < 0.01 for each antibiotic compared to relevant control. (B) and (C) the presented whole-section immunostaining images of BRCA2 and RAD51C in the blastocysts produced in control medium and antibiotics-added medium. The data were representative of at least 20 embryos for each treatment from three independent experiments.

## Discussion

In the present study, the culture of mouse zygotes to three different classes of antibiotics, even in the lowest concentration usually used in commercial ART media, gave rise to reduced developmental outcomes to the blastocyst stage. Analysis of the differential gene expression profile from RNA-seq analysis of single blastocyst found this to be associated with both down- or up-regulation of a range of genes. These involved genes implicated in multiple biological processes and signaling pathways. Of these, the expression of a number of genes associated with maintaining genomic integrity were adversely affected by exposure to antibiotics. Most notably, this included the breast cancer-related gene, *Brca2*, and a range of other related genes (suppl Figure1).

Several epidemiological studies have suggested a positive association between clinical use of antibiotic and the risk of breast cancer(28, 29), Rossini tumor and antibiotics metronidazole/cip but not gentamycin(30), whereas gentamicin may arrest cancer cell growth(31). The present study for the first time demonstrates association of antibiotics with the breast cancer related genes in the preimplantation embryo development in vitro. BRCA1/2 are tumor suppressor genes. BRCA1/2 mutations are embryo-lethal (32) and are associated with the lower ovarian reserve (33). BRCA2 physically and functionally interacts with P53 and RAD51 and is implicated in the regulation of cell cycle and DNA repair pathways(34). Antibiotics altered the immuno-detectable levels of BRCA2 and RAD51C in the embryo, whereas during IVF, a range of stressors associated with embryo culture increased the levels and nuclear localization of P53, resulting in reduced rates of development and poor postimplantation rates of survival (35). Conversely, embryo trophic ligands acting via a phosphatidylinositol-3-kinase/AKT/MDM2 signaling pathway repressed the level of P53 within embryos(36). We also noticed the different target genes of antibiotics involved in these pathways. Aminoglycoside antibiotics are related to the genotoxic response via *Atm* in as early as in 2-cell stage of embryo (37) and via *Brip1* that is more likely to increased risk of ovarian cancer. *Prkab2* as penicillin specifically downregulated helps sense and respond to energy demands within cells whereas *Brip1* was not affected by penicillin.

The BRCA mutation has been detected in IVF cycles by preimplantation genetic diagnosis (PGD) since 2008(38). PGD is a universal test for BRCA mutation carriers who require ART and fertility preservation (38-40). More recent evidences of BRCA mutation in oocyte aging and sperm aging (41, 42). However, the incidents of intragenic *Brac1/2* mutations between natural reproduction and ART is still lacking as ethical complexities of PGD are concerned. If the adverse effect of antibiotics on these tumour suppressor genes also occurs in human embryos created by ART it is conceivable that these procedures may influence the inheritance of such deleterious mutations and this hypothesis requires for the detailed investigation.

This study provides evidence for pervasive effects of commonly used antibiotics on the pattern of gene expression in the mouse embryo cultured in vitro. These changes occurred within many important regulatory pathways, but notably involve many genes performing critical functions and ensuring the integrity of the embryo’s genome. Given the centrality of DNA integrity to the future health of progeny resulting from ART procedures a detailed understanding of the potential adverse effect of this media component is an important priority for future research.

## Supporting information

Han_Manuscript Suppl Documents

## Acknowledgements

We thank Nanjing Your Bio-tech Development Ltd. Co (Jiangbei New District, Nanjing, Jiangsu Province, China) for generous donating all embryonic culture media; KSOM and HEPES-HTF.

This work was supported by grants from the National Natural Science Foundation of China awarded to X.J (81471458), and the Australian National Health and Medical Research Council (NHMRC) awarded to C.O.

## Author contribution statement

Conception and design of experiments: X.H, C.O, and X. Jin. Experiments performed by: Q.H, X.J, Y.L, L.C, J.S, W.N, W.L and X.Jin. Data analysis performed by: Q.H, X.H and X.Jin. Reagents, materials, analysis tool contribution: Y.L, J.L, X.H and X.J. Manuscript written by: C.O and X.Jin. All authors agreed to the publication.

## Figures and legends

**Suppl Figure 1.**
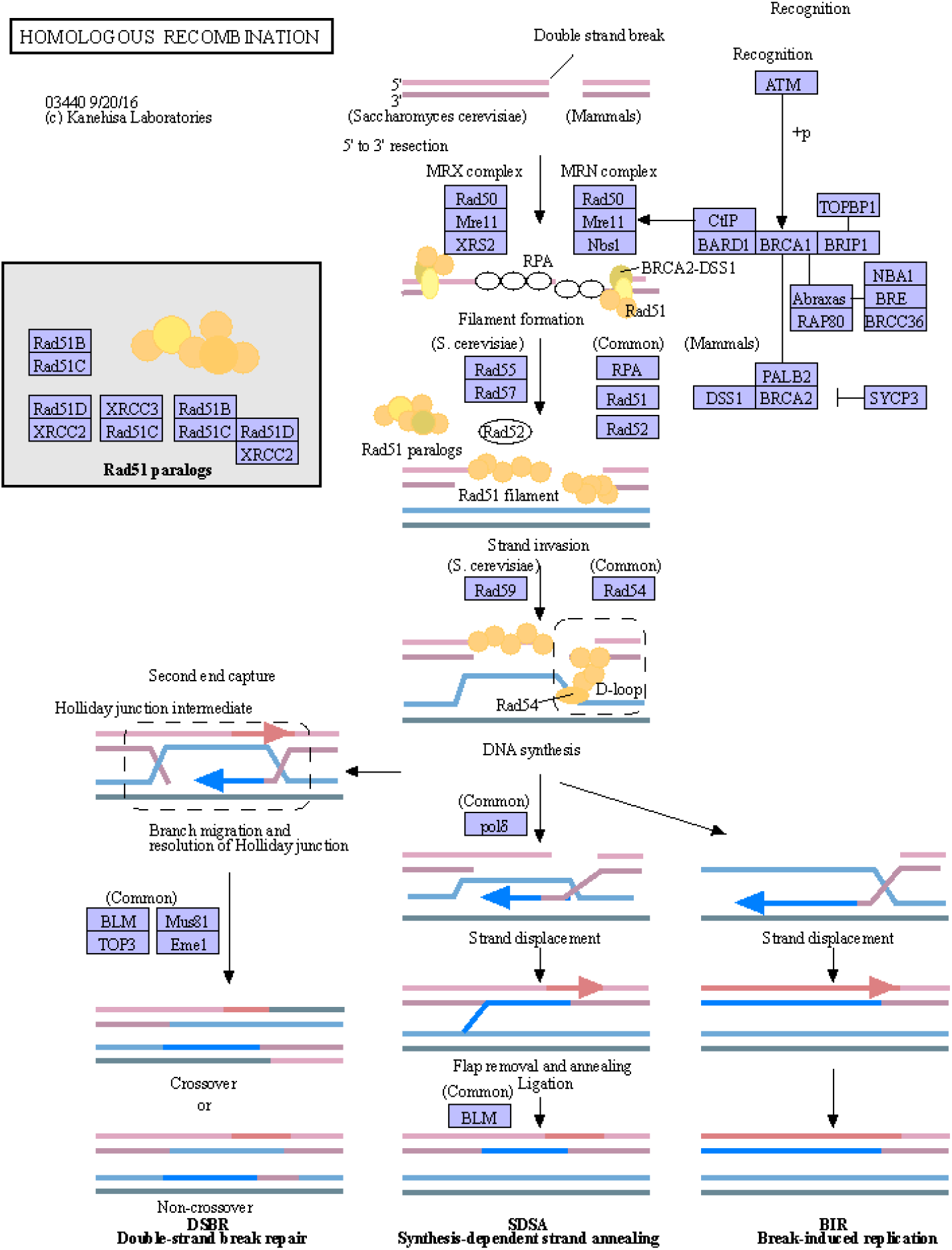
KEGG pathway for Homologous recombination (HR) in eukaryotic with relevant DE transcripts. The background diagram was adapted from Kanehisa labs (KEGG database 2018). The diagram showed that RecA/Rad51 family proteins play a central role. Brca2, which was down regulated by antibiotics, maintained genome integrity, at least in part, through HR. https://www.genome.jp/kegg-bin/show_pathway?map=ko03440&show_description=show Homologous recombination (HR) is essential for the accurate repair of DNA double-strand breaks (DSBs), potentially lethal lesions. HR takes place in the late S-G2 phase of the cell cycle and involves the generation of a single-stranded region of DNA, followed by strand invasion, formation of a Holliday junction, DNA synthesis using the intact strand as a template, branch migration and resolution. It is investigated that RecA/Rad51 family proteins play a central role. The breast cancer susceptibility protein Brca2 and the RecQ helicase BLM (Bloom syndrome mutated) are tumor suppressors that maintain genome integrity, at least in part, through HR.

