## Supplementary material for "The addition of antibiotics to embryo culture media caused altered expression of genes in pathways governing DNA integrity in mouse blastocysts": Han_Manuscript Suppl Documents: Suppl Table 1 Quality control of cNDA library and RNA-Seq data.docx

| Suppl Table 1 Quality control of cNDA library and RNA-Seq data | | | | | | | | | | |
| --- | --- | --- | --- | --- | --- | --- | --- | --- | --- | --- |
| **Samples** | **cDNA library concentration** | **Reads No** | **Reads Alignment** | **duplication** | **Transcript_ratio** | **Gene_ratio** | **5'bias** | **3'bias** | **exonic_reads_ratio** | **rRNA_ratio** |
| Control_Rep1 | 20.20 | 23136737 | 89.20% | 58.20% | 16.478% (22690/137701) | 4.111% (982/23886) | 0.49 | 0.49 | 35,069,118 (95.51%) | 0.000014% (7/50699416) |
| Control_Rep2 | 24.00 | 22949640 | 92.50% | 56.00% | 20.627% (28404/137701) | 3.718% (888/23886) | 0.55 | 0.50 | 36,674,914 (96.5%) | 0.000002% (1/50302520) |
| Control_Rep3 | 14.50 | 26437070 | 85.20% | 62.50% | 14.470% (19926/137701) | 4.626% (1105/23886) | 0.33 | 0.51 | 36,116,733 (92.54%) | 0.000000% (0/59788381) |
| **Average** | **19.57** | **24174482.33** | **88.97%** | **58.90%** | **17.19%（23673）** | **4.15% (992）** | **0.46** | **0.50** | **94.85%** |  |
| Gentamicin_Rep1 | 22.20 | 23933814 | 93.12% | 48.20% | 23.319% (32110/137701) | 3.789% (905/23886) | 0.53 | 0.49 | 38,768,609 (96.63%) | 0.000004% (2/52351632) |
| Gentamicin_Rep2 | 21.50 | 22292243 | 92.57% | 47.60% | 23.388% (32206/137701) | 3.780% (903/23886) | 0.54 | 0.48 | 36,244,797 (96.92%) | 0.000000% (0/48495369) |
| Gentamicin_Rep3 | 19.30 | 23913416 | 93.13% | 48.90% | 20.218% (27840/137701) | 3.584% (856/23886) | 0.56 | 0.53 | 38,717,388 (96.61%) | 0.000002%  (1/52319859) |
| **Average** | **21.00** | **23379824.33** | **92.94%** | **48.23%** | **22.31%（30719）** | **3.72% (888)** | **0.54** | **0.50** | **96.72%** |  |
| Streptomycin_Rep1 | 23.40 | 24572891 | 92.48% | 49.50% | 23.508% (32371/137701) | 4.505% (1076/23886) | 0.53 | 0.47 | 39,208,736 (96.17%) | 0.000006%  (3/53995110) |
| Streptomycin_Rep2 | 25.20 | 24273798 | 92.31% | 54.00% | 23.118% (31834/137701) | 4.132% (987/23886) | 0.54 | 0.48 | 38,895,950 (96.57%) | 0.000000% (0/53221631) |
| Streptomycin_Rep3 | 21.10 | 24726598 | 93.70% | 46.20% | 20.853% (28715/137701) | 3.797% (907/23886) | 0.57 | 0.50 | 40,293,110 (96.91%) | 0.000000% (0/54401112) |
| **Average** | **23.23** | **24524429** | **92.83%** | **49.90%** | **22.49%（30973）** | **4.14% (990)** | **0.55** | **0.48** | **96.55%** |  |
| Penicillin_Rep1 | 23.90 | 23263376 | 92.18% | 54.80% | 21.335% (29379/137701) | 3.986% (952/23886) | 0.60 | 0.52 | 36,997,187 (96.29%) | 0.000000% (0/51114713) |
| Penicillin_Rep2 | 21.40 | 23248427 | 91.81% | 55.60% | 19.970% (27499/137701) | 3.873% (925/23886) | 0.51 | 0.49 | 36,929,569 (96.17%) | 0.000000% (0/50725052) |
| Penicillin_Rep3 | 19.50 | 24936262 | 87.98% | 60.50% | 16.047% (22097/137701) | 4.245% (1014/23886) | 0.31 | 0.49 | 36,300,642 (94.37%) | 0.000000% (0/55325189) |
| **Average** | **21.60** | **23816021.67** | **90.66%** | **56.97%** | **19.12%（26325）** | **4.03% (964)** | **0.47** | **0.50** | **95.61%** |  |
| No-RT Rep1 | 0.42 | 2417794 | 29.95% | 23.60% | 1.049% (1445/137755) | 1.316% (594/45122) | 0.00 | 0.00 | 70,320 (10.01%) | 0.000000% (0/2467709) |
| No-RT Rep2 | 0.32 | 1615115 | 26.25% | 25.30% | 1.108% (1527/137755) | 1.270% (573/45122) | 0.00 | 0.00 | 58,444 (14.42%) | 0.000000% (0/1653256) |
| No-RT Rep3 | 1.89 | 2556848 | 42.81% | 16.50% | 1.378% (1898/137755) | 1.474% (665/45122) | 0.00 | 0.00 | 68,379 (6.45%) | 0.000000% (0/2623216) |
| **Average** | **0.88** | **2196585.667** | **33.00%** | **21.80%** | **1.18%（1623）** | **1.35% (611)** | **0.00** | **0.00** | **10.29%** |  |
