## Supplementary material for "The addition of antibiotics to embryo culture media caused altered expression of genes in pathways governing DNA integrity in mouse blastocysts": Han_Manuscript Suppl Documents: Suppl Table 2 .docx

|  | Gentamycin | Streptomicin | Penicillin |
| --- | --- | --- | --- |
| Biological Process | 1. DNA metabolic process (GO:0006259) 2. DNA recombination (GO:0006310) 3. cellular response to DNA damage stimulus (GO:0006974) 4. mitotic cell cycle phase transition (GO:0044772) 5. amyloid precursor protein catabolic process (GO:0042987) 6. DNA repair (GO:0006281) 7. regulation of cell division (GO:0051302) 8. amyloid precursor protein metabolic process (GO:0042982) 9. response to ionizing radiation (GO:0010212) 10. cellular protein modification process (GO:0006464) | 1. mitotic cell cycle phase transition (GO:0044772) 2. DNA metabolic process (GO:0006259) 3. cell cycle G2/M phase transition (GO:0044839) 4. cellular response to DNA damage stimulus (GO:0006974) 5. G2/M transition of mitotic cell cycle (GO:0000086) 6. cellular protein modification process (GO:0006464) 7. regulation of cell division (GO:0051302) 8. cell cycle G1/S phase transition (GO:0044843) 9. G1/S transition of mitotic cell cycle (GO:0000082) 10. DNA repair (GO:0006281) | 1. regulation of RNA splicing (GO:0043484) 2. phosphatidic acid biosynthetic process (GO:0006654) 3. phosphatidic acid metabolic process (GO:0046473) 4. mitotic cell cycle phase transition (GO:0044772) 5. DNA recombination (GO:0006310) 6. cell cycle G2/M phase transition (GO:0044839) 7. regulation of cholesterol metabolic process (GO:0090181) 8. vesicle transport along microtubule (GO:0047496) 9. DNA metabolic process (GO:0006259) 10. G2/M transition of mitotic cell cycle (GO:0000086) |
| List of the genes for “DNA metabolic process (GO:0006259)” | DCLRE1B;EXD2;PIF1;RTEL1;BLM;SMARCB1;HMGB3;RBPJ;BRCA2;PTMS;UNG;ALKBH3;ENDOV;BRIP1;TATDN1;RECQL5;RNASEH1;ORC6;ERI2;CHEK1;RAD54L;NT5M;MEN1;CTC1;HELLS;XRCC6;STN1;LIG1;NCOA6;RMI1;DONSON;CDC7;LIG3;CHD1L;UBE2A;DDB2;HINFP;RAD52;CDC34;RAD51C;ERCC1;CDK2;MCM4;ATM;MCM5;NUP98;RAD9B;RAD9A | DCLRE1B;MUS81;PIF1;RTEL1;BLM;SMARCB1;UHRF2;NUDT1;HMGB3;RBPJ;BRCA2;PTMS;ALKBH3;BRIP1;TATDN1;RECQL5;RNASEH1;ORC6;ERI2;CHEK1;RAD54L;NT5M;MEN1;CTC1;HELLS;XRCC6;LIG1;DONSON;CDC7;LIG3;CHD1L;UBE2A;DDB2;HINFP;RAD52;CDC34;RAD51C;ERCC1;CDK2;POLE3;KCTD13;MCM4;ATM;MCM5;NUP98;RAD9B;RAD9A | MUS81;RTEL1;BLM;SMARCB1;HMGB3;RBPJ;BRCA2;ALKBH3;ENDOV;TATDN1;RECQL5;RNASEH1;UBB;ERI2;CHEK1;RAD54L;NT5M;CTC1;HELLS;NCOA6;LIG3;CHD1L;UBE2A;RAD52;CDC34;RAD51C;ERCC1;KCTD13;MCM4;MCM5;RAD9B;RAD9A |
| List of the genes for “mitotic cell cycle phase transition (GO:0044772)” | BRSK2;CUL5;CCNH;CETN2;ARPP19;PHF8;CDC23;ORC6;HAUS8;PPP2R1A;MYC;CEP70;ENSA;RCC1;RANBP2;PLK3;NDE1;CDC7;HAUS6;MASTL;HINFP;CDC25B;DNM2;CDC34;CCNE2;CDK4;AKAP9;CDK2;TACC3;MCM4;MCM5;PAFAH1B1 | BRSK2;CUL5;CCNH;DCTN1;CETN2;ARPP19;CDC23;ORC6;HAUS8;PPP2R1A;PPME1;MYC;CEP70;ENSA;RCC1;RANBP2;PLK3;NDE1;CDC7;HAUS6;MASTL;HINFP;CDC25B;DNM2;CDC34;CCNE2;CDK4;AKAP9;CDK2;POLE3;PPP2R2D;TACC3;CHMP2A;MCM4;MCM5;CRLF3;CDK14;OPTN;PAFAH1B1 | BRSK2;CETN2;ARPP19;PHF8;CDC23;UBB;HAUS8;PPP2R1A;ENSA;RCC1;RANBP2;PLK3;ODF2;NDE1;MASTL;CDC25B;DNM2;CDC34;CCNE2;CENPJ;PPP2R2D;CHMP2A;MCM4;MCM5;CDK14;OPTN;PAFAH1B1 |

Suppl Table 2 Antibiotics on p-value-ranked Top10 GO enriched terms of down-regulated Biological Process
