## Supplementary material for "The addition of antibiotics to embryo culture media caused altered expression of genes in pathways governing DNA integrity in mouse blastocysts": Han_Manuscript Suppl Documents: Suppl Table 3.docx

Suppl Table 3 Antibiotics on p-value-ranked top 10 enriched terms of down-regulated KEGG pathways

|  | Gentamicin | Streptomycin | Penicillin |
| --- | --- | --- | --- |
| KEGG pathways | 1. Cell cycle 2. **Mannose type O-glycan biosynthesis** 3. RNA transport 4. Glycerophospholipid metabolism 5. **Homologous recombination** 6. Protein processing in endoplasmic reticulum 7. Fanconi anemia pathway 8. Tight junction 9. Ubiquitin mediated proteolysis 10. Lysine degradation | 1. Cell cycle 2. Tight junction 3. **Homologous recombination** 4. RNA transport 5. Ubiquitin mediated proteolysis 6. **Mannose type O-glycan biosynthesis** 7. Salmonella infection 8. p53 signaling pathway 9. Glycerolipid metabolism 10. Fanconi anemia pathway | 1. Glycerophospholipid metabolism 2. Choline metabolism in cancer 3. **Homologous recombination** 4. Glycerolipid metabolism 5. Fc gamma R-mediated phagocytosis 6. Sphingolipid metabolism 7. **Mannose type O-glycan biosynthesis** 8. Salmonella infection 9. Ether lipid metabolism 10. Other glycan degradation |
| List of the genes for “Homologous recombination” | RAD52;BLM;BRIP1;RAD51C;RAD54L;**ATM**;BRCC3;BRCA2 | MUS81;RAD52;BLM;BRIP1;RAD51C;RAD54L;**ATM**;BRCC3;BRCA2 | MUS81;RAD52;BLM;RAD51C;**UIMC1**;RAD54L;BRCC3;BRCA2 |
| List of the genes for “Mannose type O-glycan biosynthesis” | B3GALNT2;B4GALT3;POMT1;RXYLT1;POMGNT2;POMGNT1;LARGE2 | B3GALNT2;B4GALT3;POMT1;RXYLT1;POMGNT1;LARGE2 | B3GALNT2;POMT1;POMGNT2;POMGNT1;LARGE2 |
