## Supplementary material for "The addition of antibiotics to embryo culture media caused altered expression of genes in pathways governing DNA integrity in mouse blastocysts": Han_Manuscript Suppl Documents: Suppl Table 4.docx

| **Gentamycin** | | **Streptomycin** | | **Penicillin** | |
| --- | --- | --- | --- | --- | --- |
| **BP** | **Genes** | **BP** | **Genes** | **BP** | **Genes** |
| regulation of signal transduction by p53 class mediator (GO:1901796) | PLK3;BLM;MEAF6;EHMT2;TAF9;NOC2L;HIPK1;AURKB;TPX2;BRIP1;CHEK1;CDK2;ATM;TAF6;KMT5A;RBBP7;RAD9B;TAF4;RAD9A | regulation of signal transduction by p53 class mediator (GO:1901796) | PLK3;BLM;MEAF6;EHMT2;TAF9;NOC2L;HIPK1;AURKB;TPX2;BRIP1;CHEK1;CDK2;ATM;TAF6;KMT5A;RBBP7;RAD9B;TAF4;RAD9A | regulation of signal transduction by p53 class mediator (GO:1901796) | PLK3;PRKAB2;BLM;MEAF6;TAF9;NOC2L;HIPK1;AURKB;TPX2;UBB;CHEK1;TAF6;KMT5A;RBBP7;RAD9B;TAF4;RAD9A |
| DNA damage response, signal transduction by p53 class mediator (GO:0030330) | RBL2;PLK3;MUC1;TFDP1;PAXIP1;USP10;PCBP4;CDK2;ATM;RPL26 |  |  | regulation of intrinsic apoptotic signaling pathway in response to DNA damage by p53 class mediator (GO:1902165) | MUC1;TAF9;RPL26 |
| DNA damage response, signal transduction by p53 class mediator resulting in cell cycle arrest (GO:0006977) | RBL2;PLK3;MUC1;TFDP1;PCBP4;CDK2;ATM;RPL26 |  |  | DNA damage response, signal transduction by p53 class mediator resulting in cell cycle arrest (GO:0006977) | RBL2;CNOT4;PLK3;MUC1;UBB;PCBP4;CENPJ;RPL26 |

Suppl 4 Antibiotics caused down-regulation of *Trp53*-dependent BP (P < 0.05)
