## Supplementary material for "The addition of antibiotics to embryo culture media caused altered expression of genes in pathways governing DNA integrity in mouse blastocysts": Han_Manuscript Suppl Documents: Suppl Tables .docx

Table 1 Antibiotics on p-value-ranked GO enriched terms of Biological Process

|  | Down regulated | | Up-regulated | |
| --- | --- | --- | --- | --- |
|  | Adjusted p-Value < 0.05 | Top10 p<0.05 | Adjusted p-Value < 0.05 | Top10 p<0.05 |
| Gentamycin | 1 DNA metabolic process (GO:0006259)  2. DNA recombination (GO:0006310)  3. cellular response to DNA damage stimulus (GO:0006974)  4. mitotic cell cycle phase transition (GO:0044772)  5. amyloid precursor protein catabolic process (GO:0042987)  6. DNA repair (GO:0006281)  7. regulation of cell division (GO:0051302) | DNA metabolic process (GO:0006259)  DNA recombination (GO:0006310)  cellular response to DNA damage stimulus (GO:0006974)  mitotic cell cycle phase transition (GO:0044772)  amyloid precursor protein catabolic process (GO:0042987)  DNA repair (GO:0006281)  regulation of cell division (GO:0051302)  amyloid precursor protein metabolic process (GO:0042982)  response to ionizing radiation (GO:0010212)  cellular protein modification process (GO:0006464) | 1. positive regulation of protein localization to membrane (GO:1905477) | positive regulation of protein localization to membrane (GO:1905477)  regulation of protein insertion into mitochondrial membrane involved in apoptotic signaling pathway (GO:1900739)  positive regulation of protein insertion into mitochondrial membrane involved in apoptotic signaling pathway (GO:1900740)  positive regulation of mitochondrial outer membrane permeabilization involved in apoptotic signaling pathway (GO:1901030)  regulation of sodium ion transmembrane transporter activity (GO:2000649)  positive regulation of sodium ion transmembrane transport (GO:1902307)  phosphorylation (GO:0016310)  regulation of ion homeostasis (GO:2000021)  nucleic acid metabolic process (GO:0090304)  protein phosphorylation (GO:0006468) |
| Streptomycin (adjusted p-Value) | 1. mitotic cell cycle phase transition (GO:0044772)  2. DNA metabolic process (GO:0006259)  3. cell cycle G2/M phase transition (GO:0044839)  4. cellular response to DNA damage stimulus (GO:0006974)  5. G2/M transition of mitotic cell cycle (GO:0000086)  6. cellular protein modification process (GO:0006464)  regulation of cell division (GO:0051302)  7. cell cycle G1/S phase transition (GO:0044843)  8. G1/S transition of mitotic cell cycle (GO:0000082)  9. DNA repair (GO:0006281) | mitotic cell cycle phase transition (GO:0044772)  DNA metabolic process (GO:0006259)  cell cycle G2/M phase transition (GO:0044839)  cellular response to DNA damage stimulus (GO:0006974)  G2/M transition of mitotic cell cycle (GO:0000086)  cellular protein modification process (GO:0006464)  regulation of cell division (GO:0051302)  cell cycle G1/S phase transition (GO:0044843)  G1/S transition of mitotic cell cycle (GO:0000082)  DNA repair (GO:0006281) | - | regulation of defense response to virus (GO:0050688)  regulation of spindle organization (GO:0090224)  microtubule nucleation by microtubule organizing center (GO:0051418)  mitotic G2/M transition checkpoint (GO:0044818)  negative regulation of defense response to virus (GO:0050687)  regulation of cardiac muscle cell action potential (GO:0098901)  transcription elongation from RNA polymerase I promoter (GO:0006362)  mitotic spindle assembly (GO:0090307)  chromatin remodeling (GO:0006338)  cytoplasmic microtubule organization (GO:0031122) |
| Penicillin (p- Value) | - | regulation of RNA splicing (GO:0043484)  phosphatidic acid biosynthetic process (GO:0006654)  phosphatidic acid metabolic process (GO:0046473)  mitotic cell cycle phase transition (GO:0044772)  DNA recombination (GO:0006310)  cell cycle G2/M phase transition (GO:0044839)  regulation of cholesterol metabolic process (GO:0090181)  vesicle transport along microtubule (GO:0047496)  DNA metabolic process (GO:0006259)  G2/M transition of mitotic cell cycle (GO:0000086)  regulation of intracellular signal transduction (GO:1902531)  sphingomyelin metabolic process (GO:0006684)  intra-S DNA damage checkpoint (GO:0031573) | - | regulation of mRNA splicing, via spliceosome (GO:0048024)  RNA processing (GO:0006396)  protein ubiquitination involved in ubiquitin-dependent protein catabolic process (GO:0042787)  RNA metabolic process (GO:0016070)  peptidyl-serine autophosphorylation (GO:0036289)  mRNA splicing, via spliceosome (GO:0000398)  regulation of alternative mRNA splicing, via spliceosome (GO:0000381)  RNA splicing, via transesterification reactions with bulged adenosine as nucleophile (GO:0000377)  regulation of cellular component organization (GO:0051128)  mRNA processing (GO:0006397) |

Table 2 Antibiotics on p-value-ranked Enriched terms of KEGG (2019 mouse)

|  | Down regulated | | Up-regulated | |
| --- | --- | --- | --- | --- |
|  | Adjust p-Value < 0.05 | Top10 p<0.05 | Adjust p-Value < 0.05 | Top10 p<0.05 |
| Gentamycin | 1. Cell cycle | 1. Cell cycle 2. Mannose type O-glycan biosynthesis 3. RNA transport 4. Glycerophospholipid metabolism 5. Homologous recombination 6. Protein processing in endoplasmic reticulum 7. Fanconi anemia pathway 8. Tight junction 9. Ubiquitin mediated proteolysis 10. Lysine degradation | 1.Insulin signaling pathway | 1. Insulin signaling pathway 2. Hippo signaling pathway 3. Glucagon signaling pathway 4. Signaling pathways 5. regulating pluripotency of stem cells 6. ErbB signaling pathway 7. Non-small cell lung cancer 8. Gastric cancer 9. Basal transcription factors 10. Neurotrophin signaling pathway 11. Long-term potentiation |
| Streptomycin | 1. Tight junction | 1. Cell cycle; 2. Tight junction; 3. Homologous recombination; 4. RNA transport; 5. Ubiquitin mediated proteolysis; 6. Mannose type O-glycan biosynthesis; 7. Salmonella infection; 8. p53 signaling pathway; 9. Glycerolipid metabolism; 10. Fanconi anemia pathway | - | 1. Selenocompound metabolism; 2. Thermogenesis; 3. Lysine degradation; 4. Thiamine metabolism |
| Penicillin | - | 1. Glycerophospholipid metabolism 2. Choline metabolism in cancer 3. Homologous recombination 4. Glycerolipid metabolism 5. Fc gamma R-mediated phagocytosis 6. Sphingolipid metabolism 7. Mannose type O-glycan biosynthesis 8. Salmonella infection 9. Ether lipid metabolism 10. Other glycan degradation | - | 1. Spliceosome; 2. Other glycan degradation; 3. Fructose and mannose metabolism; 4. Hedgehog signaling pathway |

Table 3 Antibiotics on p-value-ranked enriched term of ClinVar

|  | Down regulated (p< 0.05) | Up-regulated (p< 0.05) |
| --- | --- | --- |
| Gentamycin | 1. fanconi anemia  2. alzheimer's disease  3. hermansky-pudlak syndrome  4. neoplasm of the breast  5. familial cancer of breast | 1. glycogen storage disease  2. ashkenazi jewish disorders  rasopathy |
| Streptomycin | 1. fanconi anemia  2. amyotrophic lateral sclerosis  3. neoplasm of the breast  4. familial cancer of breast  5. atrial septal defect  6. wilms tumor 1  7. hereditary nephrotic syndrome | 1. combined oxidative phosphorylation deficiency  2. xeroderma pigmentosum  3. mycobacterium tuberculosis, susceptibility to  4. amyotrophic lateral sclerosis |
| Penicillin | 1. hermansky-pudlak syndrome  2. charcot-marie-tooth disease, type i  3. peroxisome biogenesis disorders, zellweger syndrome spectrum  4. wilms tumor 1  5. hereditary nephrotic syndrome | 1. alpha-dystroglycan related dystrophy  2. mycobacterium tuberculosis, susceptibility to |

Table 4 Adjusted p-Value-ranked Enriched terms of Transcription factor PPIs

| Gentamycin | Down regulated adjusted p < 0.05 (48) | p < 0.05 (83) | Up-regulated adjusted p < 0.05 (2) | p < 0.05 (33) |
| --- | --- | --- | --- | --- |
|  | ESR1  BRCA1  TP53  SREBF2  SMAD2  NR3C1  ESR2  JUN  MYC  HDAC2  FOS  CHD1  USF1  CEBPA  POU5F1  POLR2A  SP1  CTNNB1  HTT  ATF2  AR  ETS1  BCL11A  RNF2  JARID2  NCOR1  NR4A1  PML  MITF  HCFC1  ELK4  KLF5  ZFPM2  CEBPB  MEF2A  RCOR3  BACH1  EPAS1  TBP  IRF8  HSF1  YY1  RARA  AHR  EP300  POLR3A  TRIM28  TCF12 | ESR1  BRCA1  TP53  SREBF2  SMAD2  NR3C1  ESR2  JUN  MYC  HDAC2  FOS  CHD1  USF1  CEBPA  POU5F1  POLR2A  SP1  CTNNB1  HTT  ATF2  AR  ETS1  BCL11A  RNF2  JARID2  NCOR1  NR4A1  PML  MITF  HCFC1  ELK4  KLF5  ZFPM2  CEBPB  MEF2A  RCOR3  BACH1  EPAS1  TBP  IRF8  HSF1  YY1  RARA  AHR  EP300  POLR3A  TRIM28  TCF12  TCF7L2  TFAP2A  KAT2A  SUZ12  MEF2C  CIITA  SMC3  ESRRA  YAP1  SRF  ZNF148  SALL1  NR1I2  RXRA  SMAD3  MYB  KDM5A  TFAP2C  BCLAF1  KDM6A  VDR  ELF1  NFKB1  EZH2  ELK1  PPARG  HEY1  FLI1  ESRRB  SMARCA4  RUNX3  FOXM1  NACC1  CCNT2  FOXP3 | FOXP3  USF1 | FOXP3  USF1  CTNNB1  POU5F1  RCOR1  TP53  E2F6  VDR  ZEB1  BRCA1  GATA4  WT1  HTT  WRNIP1  KDM6A  E2F4  CREB1  NFE2  AR  KAT2A  BHLHA15  HDAC2  RARA  ESRRB  ZNF217  GABPB1  YY1  TCF12  ESR1  MYC  RCOR3  STAT1  MAX |
| Streptomycin | HSF1  SREBF2  ZFPM2  TRIM28  YAP1  NFKB1  HTT  FOS  AR  NCOR1  POLR2A  JARID2  USF1  TBP  SMAD2  TCF12  KAT2A  CEBPB  ELK4  CEBPA  BACH1  NR4A1  CIITA  RNF2  PPARG  IRF8  AHR  MITF  EPAS1  SIRT3  RCOR3  PML  YY1  FOXP3  TFAP2A  ATF2  POLR3A  MEF2A  SRF  EP300  CCNE1  ELK1  MEF2C  ESRRB  STAT1  ESRRA  FLI1  NACC1  CCNT2  PPARGC1A  RARA |  | FOXP3  KDM5A  POLR2A  ESR1  HTT  RARA  RNF2  POU5F1  MYC  CTCF  ESR2  ATF2  WT1  BMI1  BHLHA15  NFE2  BRCA1  POU2F2  HDAC2  GABPB1  EP300  BRF2  NCOA1  TP63  REST |  |
| Penicillin | JUN  BCL11A  ESR1  ESR2  BRCA1  CIITA  TP53  NCOR1  NR3C1  HDAC2  POU5F1  BCL11B  BACH1  SREBF2  HSF1  TBP  HCFC1  ZFPM2  TRIM28  CHD1  MYC  FOXP3  FOS  SRF  YY1  SMARCA4  HTT  PML  POLR2A  YAP1  AR  ETS1  TAF1  AHR  MITF  FLI1  KLF9  SMAD2  USF1  STAT5A  EP300  BCL3  CTNNB1  E2F4  CEBPB |  | NFE2  EP300  STAT2  SMAD2  ESR2  FOXP3  ESR1  RUNX3  GLI1  TP53  SP1  FOXP2  HDAC2  NFE2L2  E2F6  NOTCH1  JUNB  EPAS1  KLF2  NR1I2  VDR  EED  JUN  ILF3  REST  SMARCB1  PPARG  GLIS2  PPARD  POU5F1  ERG  POLR2A  HTT  HIF1A  RCOR1  ESRRB  RUNX1  RNF2 |  |

KEGG	Homologous recombination	<https://www.genome.jp/kegg-bin/show_pathway?map=ko03440&show_description=show>

Homologous recombination (HR) is essential for the accurate repair of DNA double-strand breaks (DSBs), potentially lethal lesions. HR takes place in the late S-G2 phase of the cell cycle and involves the generation of a single-stranded region of DNA, followed by strand invasion, formation of a Holliday junction, DNA synthesis using the intact strand as a template, branch migration and resolution. It is investigated that RecA/Rad51 family proteins play a central role. The breast cancer susceptibility protein Brca2 and the RecQ helicase BLM (Bloom syndrome mutated) are tumor suppressors that maintain genome integrity, at least in part, through HR.

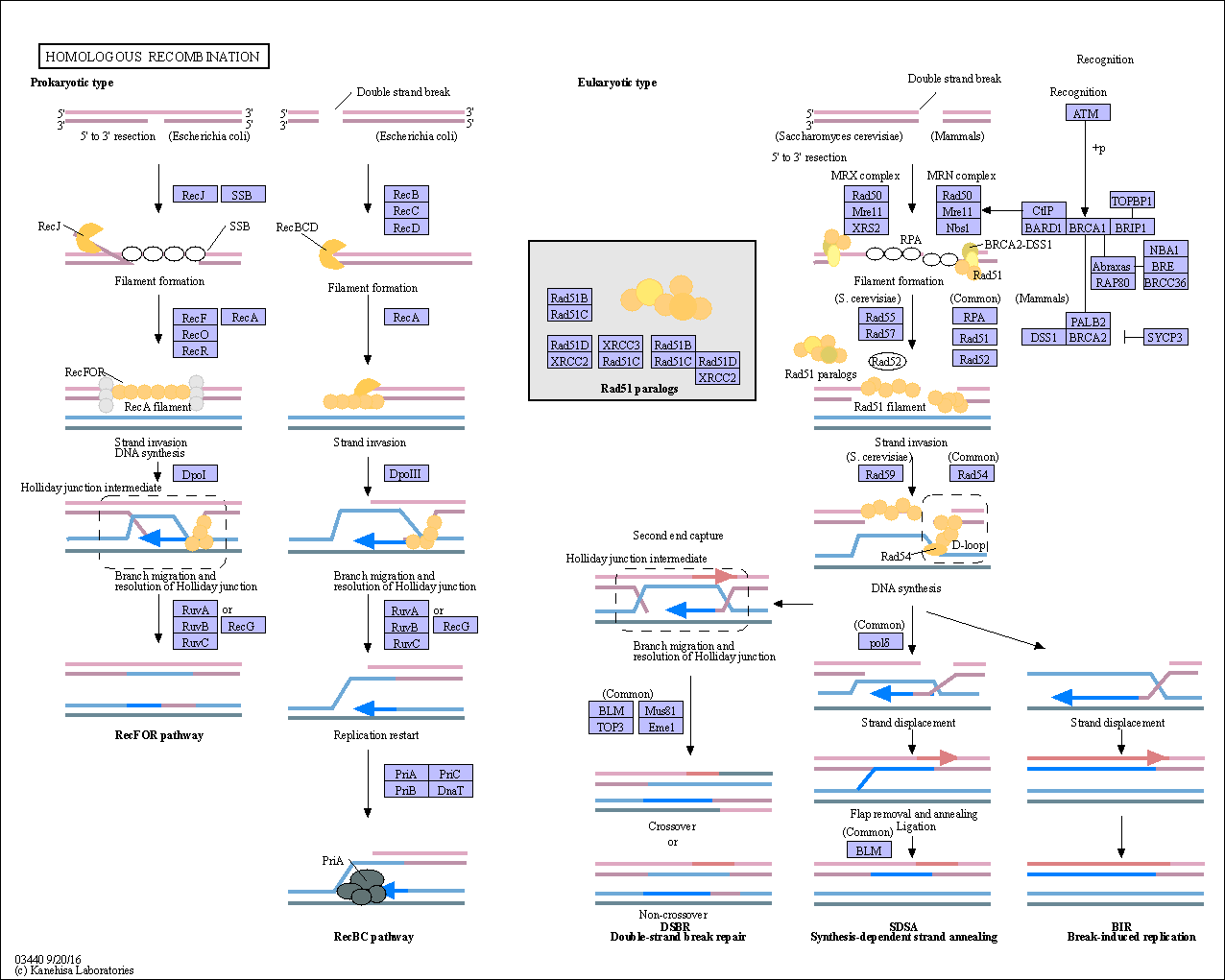
